# Embracing imperfection: machine-assisted invertebrate classification in real-world datasets

**DOI:** 10.1101/2021.09.13.460161

**Authors:** Jarrett Blair, Michael D. Weiser, Kirsten de Beurs, Michael Kaspari, Cameron Siler, Katie E. Marshall

**Affiliations:** Department of Zoology, University of British Columbia, Vancouver, BC V6T 1Z4, Canada; Department of Biology, University of Oklahoma, Norman, OK 73019-0235, USA; Sam Noble Oklahoma Museum of Natural History, University of Oklahoma, 2401 Chautauqua Ave., Norman, OK 73072-7029, USA

## Abstract

1. Despite growing concerns over the health of global invertebrate diversity, terrestrial invertebrate monitoring efforts remain poorly geographically distributed. Machine-assisted classification has been proposed as a potential solution to quickly gather large amounts of data; however, previous studies have often used unrealistic or idealized datasets to train their models.
2. In this study, we describe a practical methodology for including machine learning in ecological data acquisition pipelines. Here we train and test machine learning algorithms to classify over 56,000 bulk terrestrial invertebrate specimens from morphometric data and contextual metadata. All vouchered specimens were collected in pitfall traps by the National Ecological Observatory Network (NEON) at 27 locations across the United States in 2016. Specimens were photographed, and morphometric data was extracted as feature vectors using ImageJ. Issues stemming from inconsistent taxonomic label specificity were resolved by making classifications at the lowest identified taxonomic level (LITL). Taxa with too few specimens to be included in the training dataset were classified by the model using zero-shot classification.
3. When classifying specimens that were known and seen by our models, we reached an accuracy of 72.7% using extreme gradient boosting (XGBoost) at the LITL. Models that were trained without contextual metadata underperformed models with contextual metadata by an average of 7.2%. We also classified invertebrate taxa that were unknown to the model using zero-shot classification, with an accuracy of 39.4%, resulting in an overall accuracy of 71.5% across the entire NEON dataset.
4. The general methodology outlined here represents a realistic application of machine learning as a tool for ecological studies. Hierarchical and LITL classifications allow for flexible taxonomic specificity at the input and output layers. These methods also help address the ‘long tail’ problem of underrepresented taxa missed by machine learning models. Finally, we encourage researchers to consider more than just morphometric data when training their models, as we have shown that the inclusion of contextual metadata can provide significant improvements to accuracy.

## Introduction

Several recent studies have suggested that terrestrial invertebrates may be suffering drastic population and diversity losses (Dirzo et al., 2014; Welti, Roeder, De Beurs, Joern, & Kaspari, 2020; Wepprich, Adrion, Ries, Wiedmann, & Haddad, 2019). However, these losses are not distributed equally across the planet nor across taxonomic diversity (Guzman, Johnson, Mooers, & M’Gonigle, 2021; van Klink et al., 2020). Generalizations about trends in global invertebrate diversity and abundance require a solid data foundation, yet invertebrates remain significantly poorly-sampled relative to their diversity and abundance (Høye et al., 2021; van Klink et al., 2020). While there are many invertebrate biomonitoring programs operating around the world, the data collected from these programs are often spatiotemporally coarse and lack taxonomic diversity (Høye et al., 2021). Large-scale invertebrate biomonitoring efforts require vast funding and human resources, reducing their accessibility to developing nations and further limiting the geographic scope of biodiversity data (Karlsson, Hartop, Forshage, Jaschhof, & Ronquist, 2020; van Klink et al., 2020). One of the most time and resource intensive aspects of any invertebrate biomonitoring program relates to specimen sorting and taxonomic identification (Karlsson et al., 2020). While these processes are traditionally performed by trained taxonomists, often in coordination with parataxonomists, advances in machine learning and computer vision have made machine-assisted classification a possibility for many groups of insects (Ärje et al., 2020; Blair et al., 2020; Marques et al., 2018; Mayo & Watson, 2007). Not only does machine-assisted classification have the potential to increase the efficiency, output, and accessibility of biomonitoring programs (Peters et al., 2014; Thessen, 2016), but also, in a time when more data on invertebrate diversity and abundance is desperately needed, such an approach offers a transformative solution to the way we monitor global invertebrate diversity in a changing climate.

Excitement regarding the possibilities of machine learning for taxonomic classification has led to many publications on the topic over the last decade. However, a common theme in this literature is the use of unrealistic or idealized case studies. Specifically, many datasets used in these studies have low species richness, uniform taxonomic resolution, and no geographic/temporal component (Ärje et al., 2020; Blair et al., 2020; Joutsijoki et al., 2014). Additionally, nearly all studies that have used large ecological datasets accept that low abundance taxa must be sacrificed for the benefit of their model’s performance (Marques et al., 2018; Mayo & Watson, 2007). In most natural communities, most taxonomic diversity is comprised of low-abundance taxa (i.e. the long-tail of rank abundance curves; Preston, 1948; Verberk, 2012; Whittaker, 1965), so simply ignoring them is not feasible if machine learning tools are to be widely used in ecological studies. We note that the previously-mentioned machine learning papers still provide foundational knowledge about machine learning’s potential uses for taxonomic classification. However, significant work remains to improve the practicality of computer vision for specimen classification if it is to be integrated into ecological data acquisition pipelines.

One practical application of machine learning for ecological purposes is the web and mobile app iNaturalist (Van Horn et al., 2018). iNaturalist is a community science social network in which users can share their own biodiversity observations as well as comment on those submitted by others. In 2018, iNaturalist implemented an automated identification tool, whereby a computer vision model recommends taxonomic classifications based on user submissions (Van Horn et al., 2018). Predecessors of iNaturalist such as LeafSnap and the Merlin Bird ID applications had similarly implemented automated classifications using computer vision (Kumar et al., 2012; Van Horn et al., 2015), but iNaturalist was the first to do so with the goal of being able to classify *any* macro organism on the planet. However, as a community science focussed application, iNaturalist has limitations that make it unsuitable for other macroecological uses. For example, iNaturalist submissions tend to be biased towards charismatic taxa, which causes its automated identification tool to misidentify uncharismatic fauna such as ground beetles (Blair et al., 2020). Additionally, the current build of iNaturalist is not suited for large-scale bulk identifications that are needed for many macroecological studies. In many instances, custom built, automated classifiers are still necessary instead of off-the-shelf applications like iNaturalist.

While there have been some promising results developing models for classifying biological specimens (Blair et al., 2020; Ding & Taylor, 2016; Spiesman et al., 2021; Van Horn et al., 2018), ecological datasets pose several challenges for machine learning. For example, most ecological datasets are uneven, with relatively few taxa that are highly abundant, while many others have relatively low abundance. This causes two problems for machine learning algorithms. First, machine learning algorithms tend to be biased towards overabundant labels. This is because a model may achieve higher accuracy by simply predicting the most abundant taxa every time instead of making informed predictions (Dinga, Penninx, Veltman, Schmaal, & Marquand, 2019). This can lead to relatively ‘accurate’ but useless models that never predict low abundance taxa (Dinga et al., 2019). Second, there may not be enough data to properly train a machine learning algorithm to identify taxa in the ‘long tail’. These taxa are ‘unknown and unseen’ by the machine learning model and therefore, can never be specifically predicted by the model (Deng, Krause, Berg, & Fei-Fei, 2012). For machine learning to be a practical solution for use in ecological studies, these problems must be addressed.

Another challenge facing the classification of ecological datasets (especially large, diverse ones) is a ‘ragged edge’ of taxonomic specification: in many ecological datasets, identifications are made at varied taxonomic resolution dependent on the taxonomist’s knowledge (Jansen, Hill, Dunstan, Eléaume, & Johnson, 2018; Schmidt-Kloiber & Nijboer, 2004). For example, some specimens may be identified down to species, while others are identified to a higher taxonomic level only (e.g. family, order, etc.). This then poses the question of what a ‘correct’ classification is, and machine learning algorithms may need to weigh the benefits of accuracy versus specificity (Deng et al., 2012). Here we explored two potential ways to address this problem: lowest identified taxonomic level (LITL) models and single-level models. Single-level models assign labels to specimens and measure accuracy all at one specified taxonomic level, while LITL models assign labels and measure accuracy based on the LITL label assigned to each individual. LITL labels allow for maximum inclusion and specificity, but result in nested labels (i.e. labels belonging to related taxa at different taxonomic levels, such as Formicidae and Insecta; Figure S1). Single-level models prevent nested labels by classifying all individuals at a single taxonomic level but can result in excluded individuals that were labelled at higher taxonomic levels in the training dataset. Additionally, single-level models restrict taxonomic specificity as classification cannot be made below the specified taxonomic level. From an ecologist’s perspective, there is likely no definitive best solution as the importance of taxonomic specificity will vary depending on the use case.

Despite the challenges ecological datasets pose, they also provide many unique opportunities for improving machine learning performance. For example, they often have contextual metadata about the collection site (e.g. location, time, temperature, etc.) that can be used to inform computer vision. This contextual metadata can significantly improve accuracy, as seen in an accuracy increase of 9.1% (48.1% → 57.3%) when classifying ladybeetles (Coleoptera: Coccinellidae; Terry et al., 2020), an 11.2% increase (68.6% → 79.8%) when classifying North American birds (Berg et al., 2014), and a significant improvement when classifying marine plankton (Ellen, Graff, & Ohman, 2019). Additionally, the hierarchical structure of taxonomies can be used to perform ‘zero-shot classification’, which allows unknown and unseen classes to be classified by making predictions at the higher taxonomic rank that is known by the model (Blair et al., 2020; Deng et al., 2012). Deng et al. (2012) shows that while ‘flat’ classifiers (i.e. single-level classifiers) have zero accuracy and zero information gain when classifying unknown and unseen classes, hierarchical classifiers can often predict these classes accurately. Strategies such as these may greatly increase the practicality of incorporating computer vision methods in large scale ecology projects.

One such large scale ecology project is the National Ecological Observatory Network (NEON), which collects open access ecological data at a continental scale in the United States (Keller, Schimel, Hargrove, & Hoffman, 2008). Part of NEON’s ecological monitoring involves collecting and processing thousands of invertebrate specimens every year from arrays of pitfall traps that are collected and pooled every 14 days (Thorpe et al., 2016; Hoekman et al., 2017) (Figure S2). NEON’s current workflow for these samples is to have a local parataxonomist separate and identify all ground beetles (Family: Carabidae), while the remaining unsorted invertebrate bycatch is stored in a biorepository. While NEON’s decision to focus on carabids is understandable given the human resources it would require to properly sort the invertebrate bycatch, it also means they are sitting on a large and as-of-yet untapped source of North American terrestrial invertebrate data. This makes NEON’s invertebrate bycatch a prime target for machine assisted specimen classification so that researchers in a wide variety of disciplines can make use of this extensive data source.

Here we apply machine learning classification methods to the large terrestrial invertebrate dataset collected by NEON. Our primary goals are to address the challenges taxonomic classification poses for machine learning while also taking advantage of the benefits provided by ecological datasets. We did this by training a variety of algorithm types on several configurations of the NEON dataset to determine the optimal algorithm and dataset configuration. Using the NEON dataset, we ask: (1) How does the inclusion of contextual metadata impact model performance and what is its relative importance compared to morphometric data? And (2) how can we improve classification in the real-world conditions of low-abundance taxa and inconsistent taxonomic resolution? We predict that models trained with datasets that include contextual metadata will outperform models that only contain morphometric data. We also predict that if the presence of overrepresented groups adversely affects model performance, increasing the evenness in the training data will improve model performance, especially when measured by F1 score. Here we show that the inclusion of contextual metadata improves classification performance while unevenness in label representation is relatively unimportant. We also compare the performance and practicality of LITL vs a single-level (order) model and use zero-shot classification to overcome the ‘long tail’ problem. Taken together, we outline potential approaches for the challenges of machine-assisted classification in “real world” ecological datasets.

## Methods

### Invertebrate Bycatch Collection

We used terrestrial invertebrates collected from pitfall traps by the National Ecological Observatory Network (NEON, see Hoekman et al., 2017 for collection methodology). All specimens were collected in 2016. NEON sets the pitfall traps to collect ground beetles (Coleoptera: Carabidae) and separates them from the rest of the invertebrates, so they are not considered here (but see Blair et al., 2020).

### Imaging

We made bulk digital images of the invertebrates (mostly Arthropoda, Mollusca and Annelida) at a resolution of 729 pixels per mm^2^ against a white background (for complete methods, see Weiser, Marshall, Siler, & Kaspari, 2021; Figure S2). Using the FIJI implementation of ImageJ (Schindelin et al., 2012), for each individual organism, we extracted 21 morphological measures (e.g., major and minor axis, perimeter, image area) and 8 statistics (e.g., mean, skew, and kurtosis) for the distribution of values from each of the three RGB (Red, Green, and Blue) color layers.

### Contextual Metadata

Here, we define contextual metadata as any non-morphometric and non-taxonomic data included in the invertebrate dataset. Our contextual metadata can broadly be categorized as spatiotemporal, temperature, and precipitation data. We extracted the daily minimum and maximum temperature as well as the daily precipitation for each location from the Daymet V3 dataset. Daymet V3 provides gridded estimates of daily climatological and weather variables at 1 km^2^ spatial resolution for North America. The gridded datasets are developed using ground-based observation and statistical modeling techniques (Thornton et al., 2021). Spatiotemporal data includes longitude, latitude, and the ordinal day of the sampling event. Our temperature and precipitation data were measured daily and reported as the minimum, maximum, mean, and standard deviation during the preceding two weeks before a sampling event. All contextual metadata values were the same for all specimens collected in a given trap event. Between our morphometric data and contextual metadata, 46 descriptive variables were used as our training data (36 morphometric variables, 10 contextual variables).

### Machine Learning

#### Preparation

All machine learning methods were developed using R version 4.0.5 (R Core Team, 2021), and all code is available on GitHub (Blair, 2022). To prepare the data for machine learning training and testing, we first removed all taxa with fewer than 100 observations from the dataset (Figure 1), as these taxa contained too little data to properly train the models. We then randomly split the dataset in training and testing datasets at a ratio of 70:30. This process was repeated 10 times, resulting in 10 separate training and testing datasets (Figure S3) to ensure we had robust performance measurements for each model. Finally, we normalized the data by centering and scaling each predictor variable such that they all had a mean of 0 and standard deviation of 1. After pre-processing the data, it was ready to be used for training in a machine learning model. We contrasted five types of machine learning algorithms: K-nearest neighbours (KNN; Cover & Hart, 1967), linear discriminant analysis (LDA; Mika, Ratsch, Weston, Scholkopf, & Muller, 1999), naïve Bayes (NB; Mika et al., 1999), eXtreme Gradient Boosting (XGBoost; Chen & He, 2014), and artificial neural networks (ANN; Haykin, 2008). We chose these five algorithms as they use a wide range of machine learning techniques and increase our likelihood that at least one algorithm will reach a high classification accuracy. Our methods for optimizing these algorithms are described in Table 1.

**Table 1:**
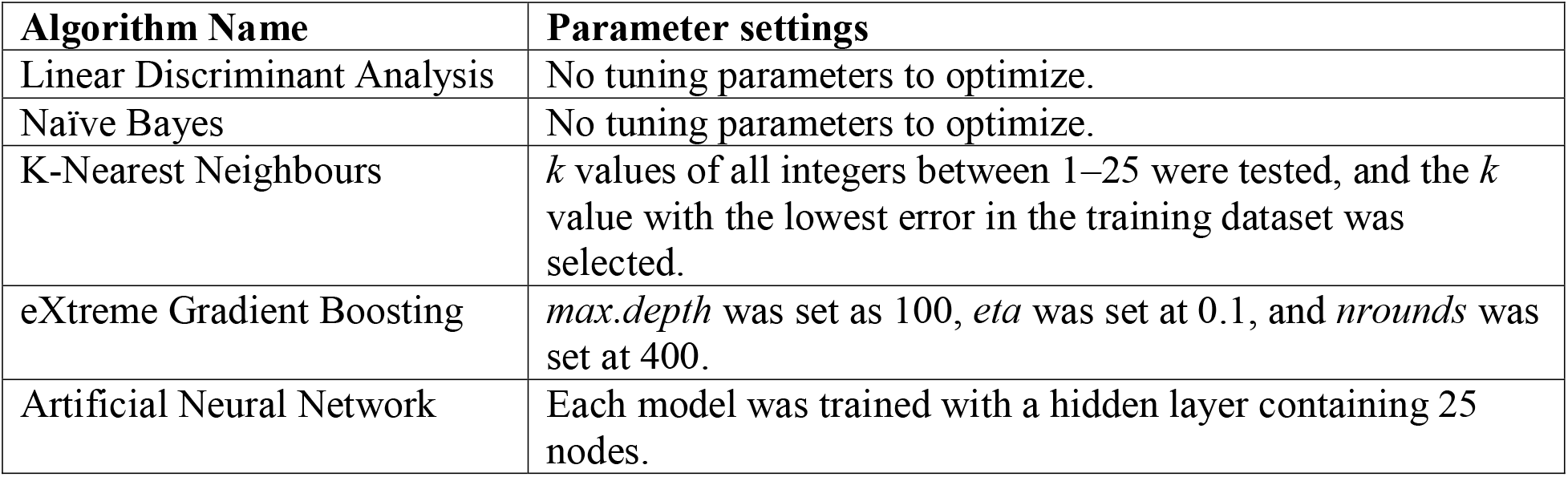
Classification algorithms and their respective parameter settings used for classifying the NEON invertebrate bycatch.

**Figure 1.**
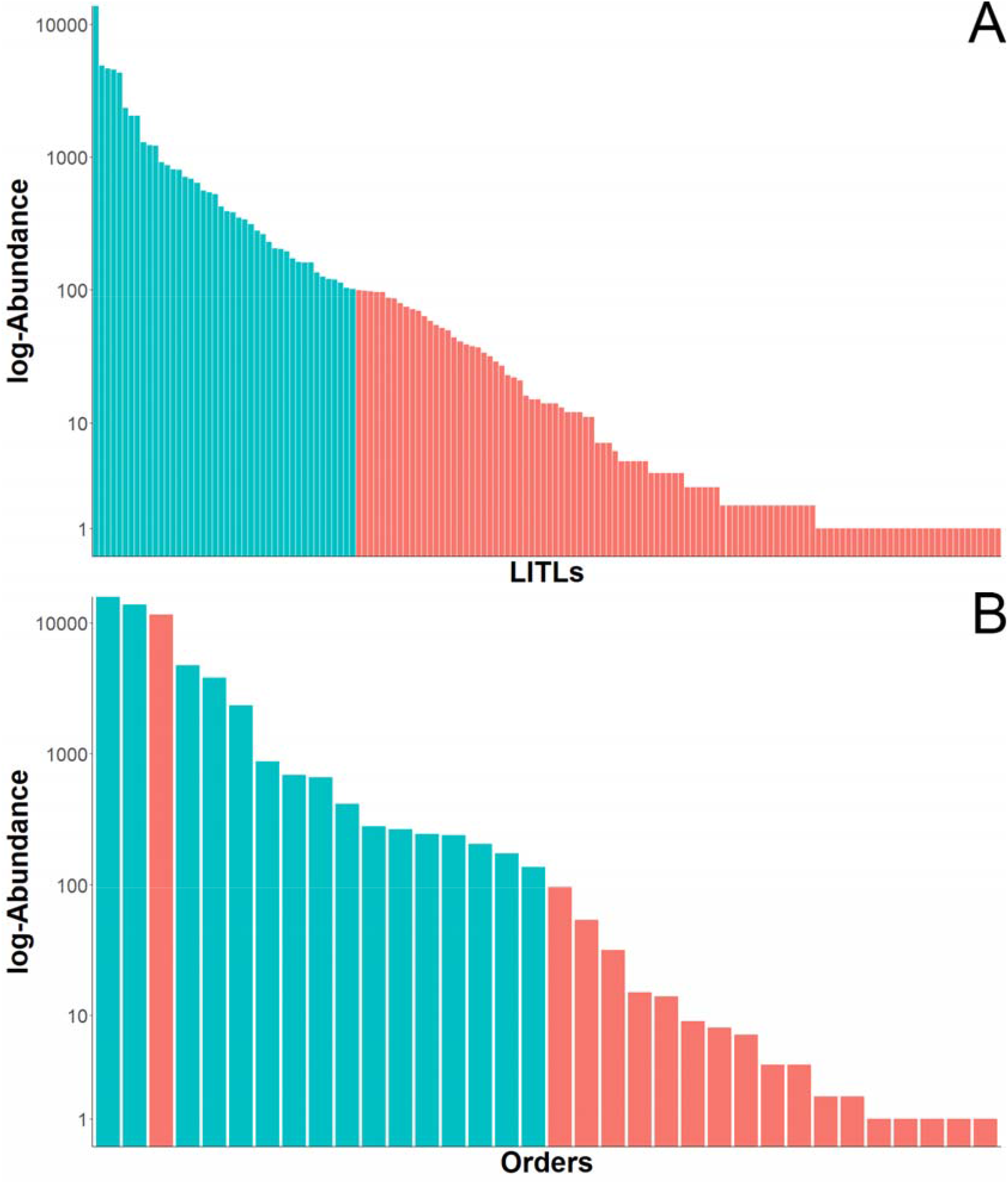
Log-scale rank abundance plots of invertebrates in the NEON dataset. (A) Each bar represents a lowest identified taxonomic label (LITL), as identified by MDW. Teal bars represent groups that had ≥100 individuals and were included in the training dataset; red bars represent groups with <100 individuals and were removed from the training dataset. (B) Each bar represents a taxonomic order of invertebrates, as identified by MDW. Teal bars represent groups that had ≥100 individuals and were included in the training dataset; red bars represent groups with <100 individuals or had a LITL above the level of order (e.g. class or phylum) and therefore were removed from the training dataset.

#### LITL & Order-level labels

Due to the uneven taxonomic resolution of our dataset (i.e. some individuals were classified to lower taxonomic levels than others), specimens in the training dataset were labelled at their lowest identified taxonomic level (LITL; Figure S1a). We treated nested taxonomic labels (e.g. Staphylinidae and Coleoptera) as non-equivalent and mutually exclusive. For example, if a specimen with a LITL label of Staphylinidae is classified as Coleoptera by a model, or vice-versa, that classification would be deemed incorrect despite Staphylinidae being a family within Coleoptera. We also created a separate training dataset in which all specimens were labelled at the order level (Figure S1b). Any specimens with an LITL label below the order level were relabelled as their corresponding order (e.g. Staphylinidae would be relabelled as Coleoptera). Conversely, any specimens with an LITL label above order level were removed from the training dataset. These datasets were pre-processed separately from the LITL datasets and included orders with 100 or more individuals in the NEON dataset. Performance of the LITL and order-level models were measured using accuracy and F1 score.

#### Contextual Metadata

To determine the effects and importance of contextual metadata, we trained and tested models of each algorithm type using LITL datasets that contained both contextual metadata and morphometric data as well as with datasets that only contained morphometric data. Differences in performance were measured as the net change in accuracy, and variable importance was measured using ‘mean decrease accuracy’ in the XGBoost model.

#### Adjusting for sampling bias

To combat the unevenness of taxonomic group representation (e.g. overabundance of ant classes), we trained and tested several configurations of binary and multiclass classifiers (Figure S4). The objective of the binary classifiers was to increase evenness among classes by removing/separating out overabundant classes. Our first binary classifier was an “adult invertebrate” classifier. When creating the invertebrate dataset, some non-adult invertebrate objects (e.g. juveniles, body parts, abiotic particles, etc.) were included. The goal of this classifier was to identify adult invertebrates and classify them as “keep” and classify everything else as “ignore”. Specimens labelled as “keep” were kept in the dataset and passed on to subsequent classifiers, while “ignore” specimens were discarded. The “keep” specimens were passed on to one of two classifiers depending on the classifier configuration. One option was for the specimens to simply be passed to a multiclass model to assign the final taxonomic classification. However, of the 44 total LITLs, 16 were related to ants (i.e. Formicidae or lower). We predicted that this may cause the model to be biased towards ant-related classifications, and designating them their own model would reduce that bias. To account for this, the specimens could be passed to a second binary classifier to split the data into “ants” and “not ants”, where “ants” are any member of the family Formicidae and “not ants” are every label not part of the “ant” or “ignore” groups. Once classified, the “ant” and “not ant” groups were passed on to their own respective multiclass models to assign the final taxonomic classification. All binary and multiclass models were trained using XGBoost with morphometric and contextual metadata.

Another method we tested to supress the effects of uneven representation was down-sampling. To do this, we randomly removed individuals from overrepresented labels in the training dataset until no label had more than 1000 specimens. Overrepresented taxa were not down-sampled from the testing dataset, as this maintained their natural occurrence frequencies and allowed for a 1:1 performance comparison with the other models. All down-sampled models were trained using XGBoost with morphometric and contextual metadata.

#### Zero-shot classification

We performed zero-shot classification by taking taxa that had too few specimens to be included in our training datasets (“unseen” taxa) and classifying them at taxonomic levels where they belonged to a common group that was included in the training dataset (“known” taxa). For example, there were only 87 specimens from the family Elateridae (click beetles) in our LITL dataset, and thus this family was not included in the training dataset. This makes the label Elateridae unknown and unseen by the models, which means that the models cannot classify this label at the family level. However, Elateridae belongs to the order Coleoptera, which is a label known by the model. This allowed us to make a zero-shot classification for Elateridae at the order level and above by classifying them as Coleoptera. When measuring zero-shot accuracy, any classification made of a label belonging to the known group would be deemed correct. For example, if “Elateridae” was classified as “Staphylinidae” instead of “Coleoptera”, it would still be considered accurate at the order level despite “Staphylinidae” being treated as mutually exclusive from “Coleoptera” in the LITL models. We note that in practical situations uncommon groups could be added to the training dataset by labelling them at their known taxonomic level, but they were left out of our training dataset so we could measure zero-shot classification performance. We also performed zero-shot classification using the order level models by classifying unincluded taxa at the class and phylum levels.

We measured the accuracy of zero-shot classifications in two ways (Table S1). In the first method, hereon known as the comprehensive method, we measured accuracy using all zero-shot specimens at every taxonomic level, regardless of if they had a known taxonomic label. For example, in the order-level models, a specimen with the LITL of “Arthropoda” would still be included when calculating the class-level zero-shot accuracy despite having zero chance of being correctly classified by the model. In the second zero-shot accuracy measurement, hereon known as the limited method, we only calculated accuracy using specimens that had a known taxonomic group. Using the limited method, specimens with the LITL of Arthropod would only be included in the accuracy calculation at the phylum-level. When measuring zero-shot accuracy, specimens labelled as any non-intact adult group (i.e. “Ignore”, “Larva”, etc.) were classified as the species level in the LITL models and at the order level in the order level models. We also used zero-shot accuracy to estimate LITL and order-level accuracy across the entire invertebrate dataset. We extrapolated the accuracies measured from the testing datasets to the training datasets to get an estimated accuracy for all ‘known’ specimens (43,025 LITL specimens and 43,390 order-level specimens). We then combined this estimated accuracy with our zero-shot accuracies to get an estimated accuracy for the entire dataset.

We measured the taxonomic specificity of the LITL and order level models by averaging the taxonomic level of each label in the training and testing datasets, as well as the first known levels for zero shot specimens. Six taxonomic levels (species, genus, family, order, class, and phylum) were used. Each taxonomic level was assigned a numerical value in ascending order (i.e. species

= 1, phylum = 6). If the taxonomic level of a label was between one of the six measured levels, its value was rounded up (e.g. superfamily labels were measured as order level). Specimens labelled as “Ignore”, “Larva”, “Nymph”, and “Juvenile” were not assigned a numerical value and were removed from the measurement.

## Results

### LITL

Our entire NEON invertebrate dataset contained 56,153 specimens with 153 LITLs and 29 taxonomic orders (Figure 1). After we removed uncommon LITL labels (<100 individuals per label), our training datasets contained between 37,913 and 37,975 specimens across 44 LITL labels while our testing datasets contained between 16,298 and 16,236 specimens (Figure 1). Our training datasets varied in size due to the random sampling method we used when splitting the datasets. The 1941 uncommon specimens we removed from the training and testing datasets were used for zero-shot classifications. Some (2 – 8 out of 10, depending on the training data) neural networks did not converge and thus did not produce a working model. Only neural networks that converged were included in performance metric calculations. The performance metrics of all other models were taken as the average across the 10 iterations.

We found models that contained metadata (i.e. location, date, temperature, and precipitation data) always performed better than their non-metadata counterparts on all metrics across all algorithms when predicting specimens down to their LITL (Figure 2). Of the models we tested, XGBoost always performed best, with an average top-1 accuracy of 72.7%, top-3 accuracy of 90.3%, and F1 score of 0.651 when including metadata. XGBoost also had the greatest accuracy increase between the metadata and no-metadata configurations with an average boost of 9.0%(63.7→ 73.7%). The average accuracy boost when we included metadata in the training dataset was 7.2% across all models. We also found the F1 score of KNN was improved the most when metadata was added (+0.160), with an average increase of 0.121 across all models. Latitude and longitude had the highest importance (mean decrease in accuracy) among metadata variables for the XGB models, ranking sixth and ninth out of 46 predictor variables respectively (Figure S5).

**Figure 2.**
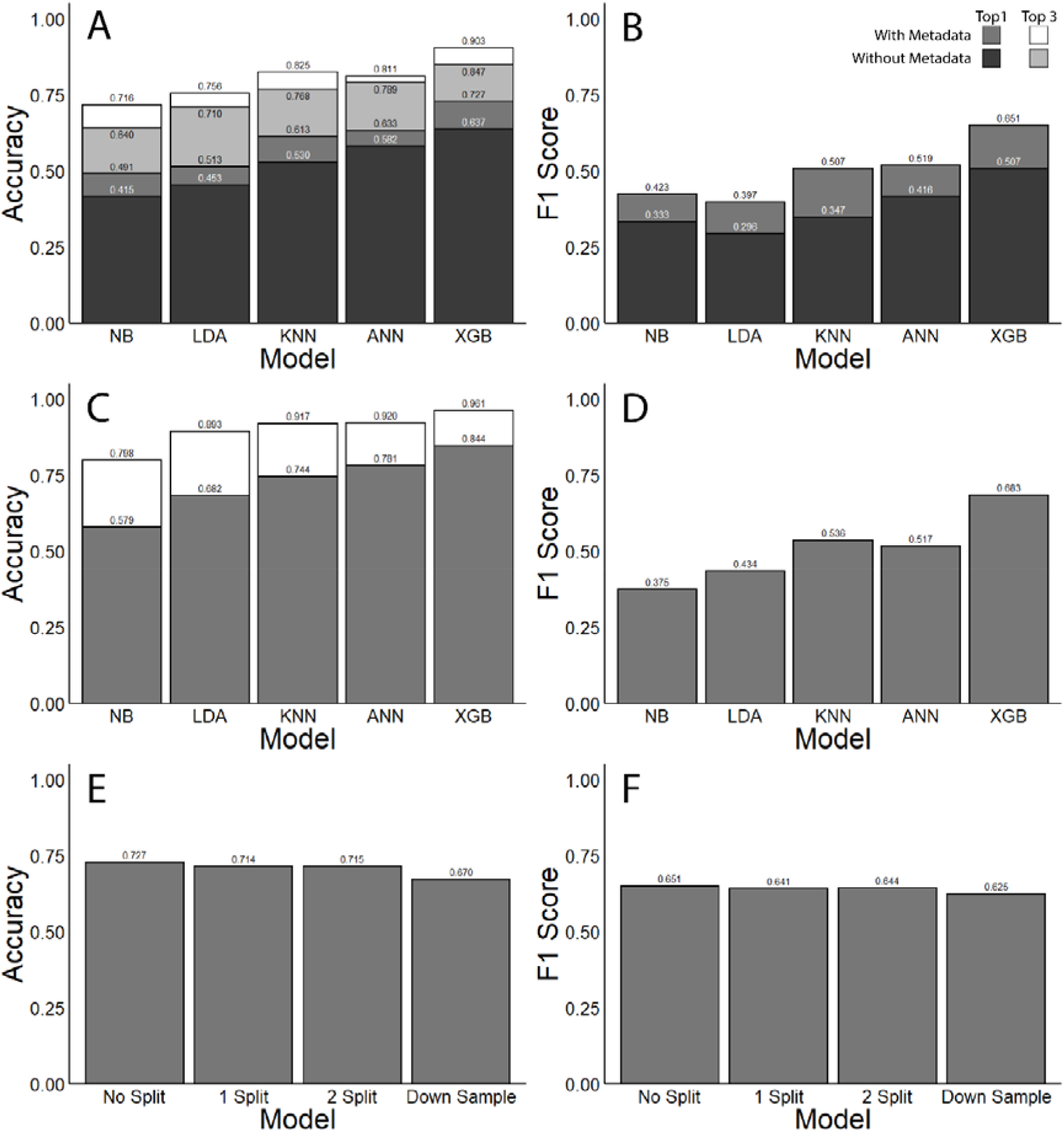
(A, B, C, D) Invertebrate bycatch classification performance metrics for models trained with and without contextual metadata All results are shown as the average across 10 models for each algorithm, except for ANN which only has one model. Top 1, Top 3, and F1 score are defined in the Glossary. (A) Top 1 and top 3 accuracy at LITL. (B) F1 score at LITL. (C) Top 1 and top 3 at order level. (D) F1 score at order level. (Naïve Bayes [Bayes]; Linear discriminant analysis [LDA]; K-Nearest Neighbours [KNN]; Artificial Neural Network [ANN]; Extreme Gradient Boosting [XGB]). (E, F) Invertebrate bycatch classification performance metrics of the extreme gradient boosting (XGBoost) model across four model configurations. All results are shown as the average across 10 XGBoost models trained with contextual metadata. ‘No splits’, ‘1 split’, and ‘2 splits’ are defined in the Glossary. ‘Down Sample’ uses no binary classifiers, but the most abundant classes in the training dataset were down-sampled to <1000 specimens. (E) Accuracy results of each XGBoost model configuration. (F) F1 score of each XGBoost model configuration.

Tactics to increase evenness in taxonomic representation (e.g. splitting ants into their own model, down sampling, removing “Ignore” groups) did not increase accuracy (Figure 2). Down sampling resulted in the lowest accuracy (67.0%) and F1 score (0.625). Splitting the dataset twice resulted in no change in accuracy but resulted in an F1 score of 0.644.

We performed zero-shot classification on unseen and unknown specimens by classifying them as known classes at higher taxonomic levels. When classified to the lowest possible level, we found zero shot classification had a top-1 accuracy of 39.4%. The zero-shot accuracy increased as the taxonomic resolution of classifications decreased, with an accuracy of 76.0% at phylum (Figure 3). When we combined regular classifications and zero-shot classifications, the average overall top-1 accuracy of the XGBoost model across the entire NEON dataset was 71.5% when classifying to the lowest possible label with metadata included. The average specificity for the LITL model was 3.7 (i.e. between family and order).

**Figure 3.**
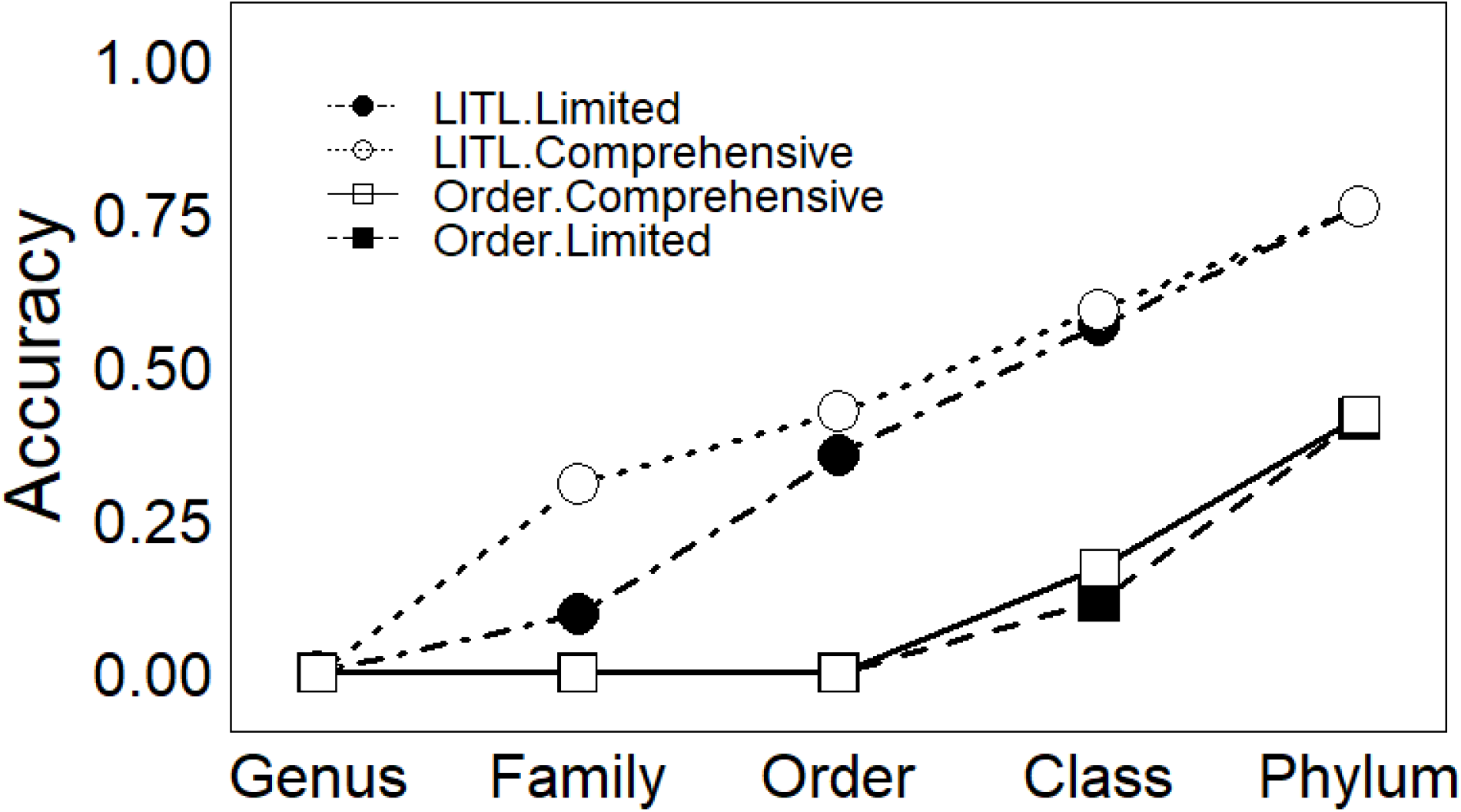
Zero-shot XGBoost accuracy plot across all tested taxonomic levels. Lowest Identified Taxonomic Level (LITL) label trained models are represented by circles, while order-level models are shown as squares. Accuracy measured using the “limited” method is shown by solid shapes, while accuracy measured using the “comprehensive” method is shown with hollow shapes (Table 2).

### Order level

After we removed uncommon orders and specimens with LITLs above order (Figure 1), our training datasets contained between 31,035 and 31,179 specimens across 16 orders (including “Ignore”, “Larva”, and “Nymph” groups) while our testing datasets contained between 13,250 and 13,393 specimens. The 11,724 specimens we removed from the training and testing datasets were used for zero-shot classifications.

**Table 2:**
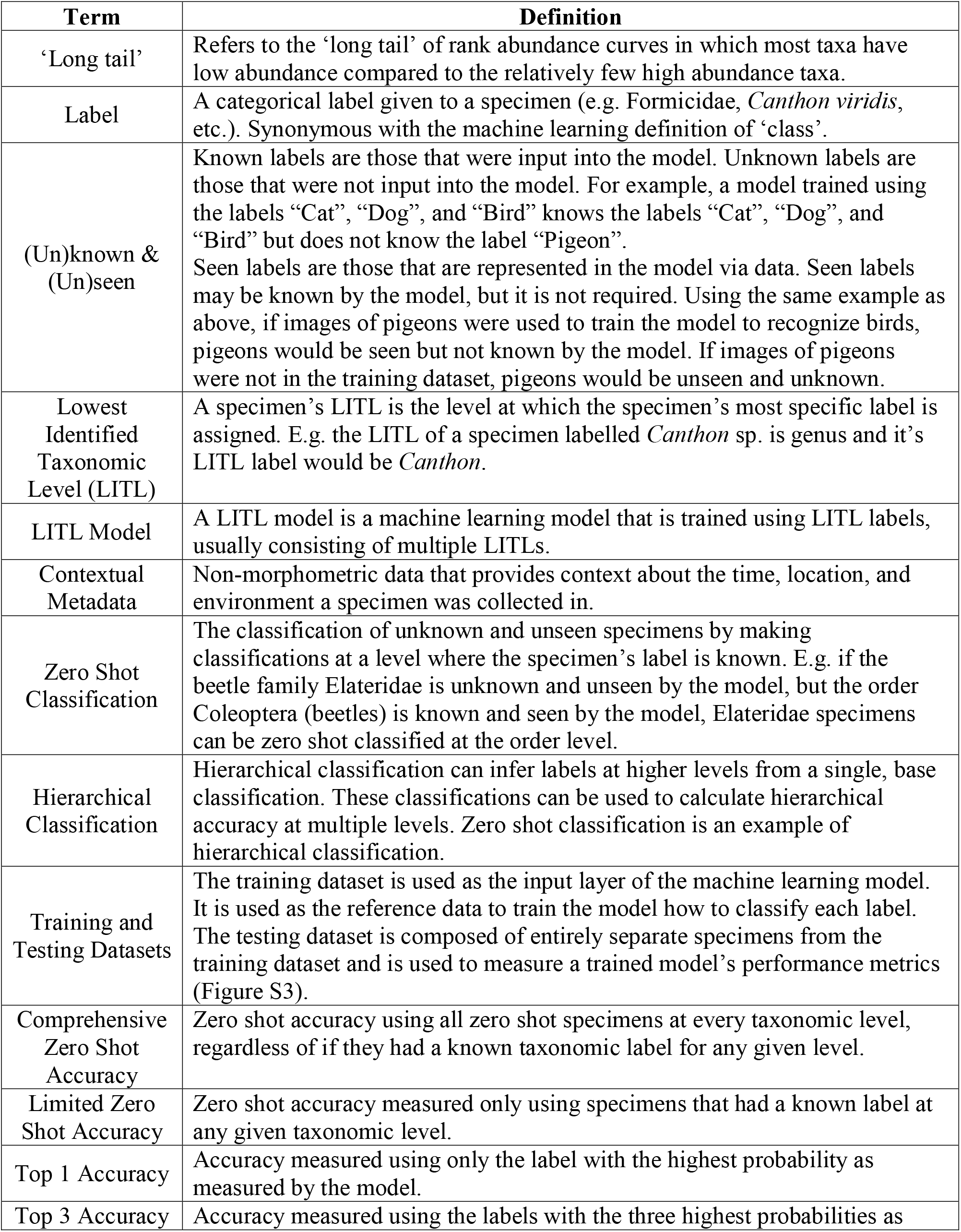
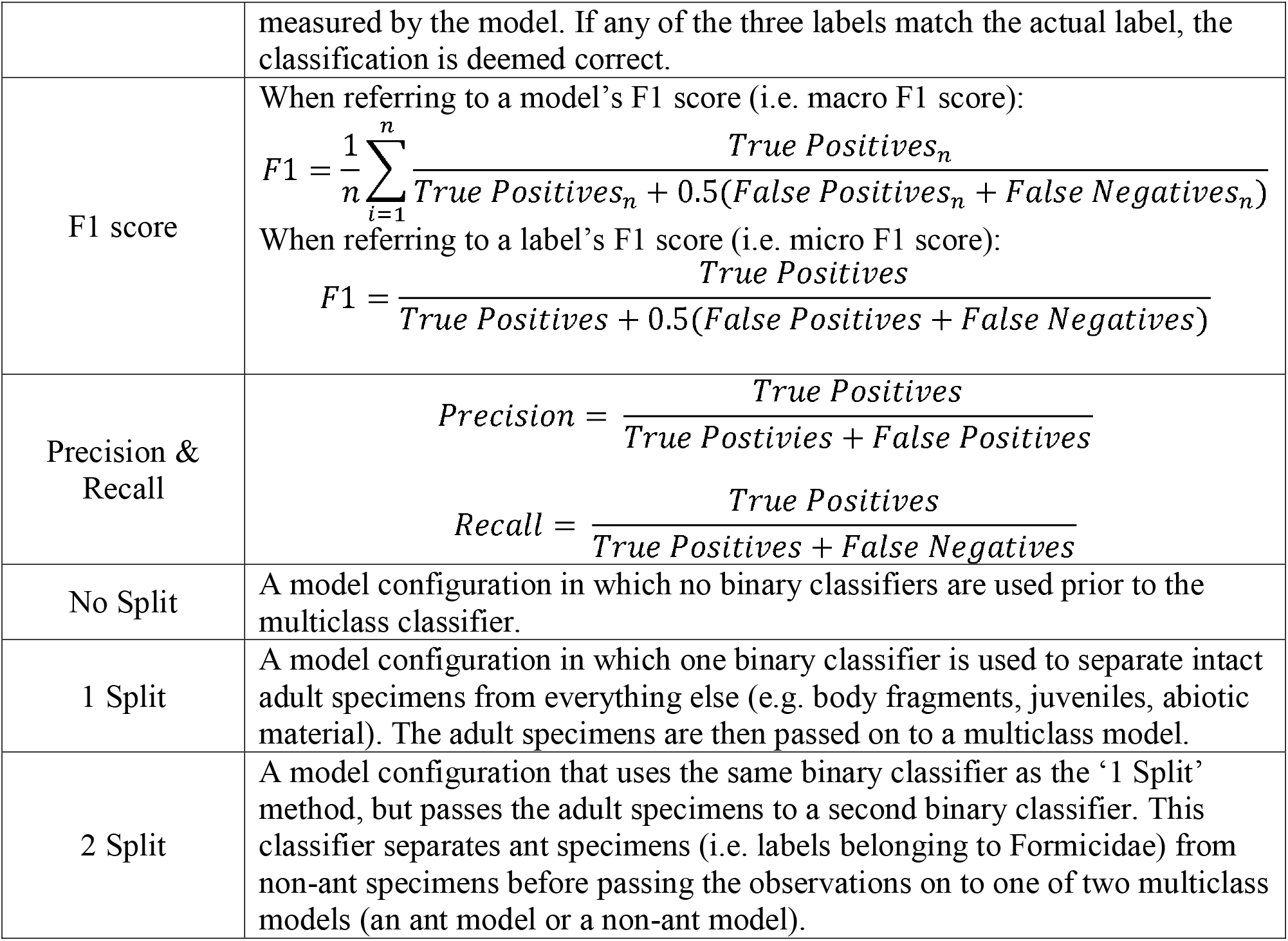
Glossary of terms.

| Term | Definition |
| --- | --- |
| ‘Long tail’ | Refers to the ‘long tail’ of rank abundance curves in which most taxa have low abundance compared to the relatively few high abundance taxa. |
| Label | A categorical label given to a specimen (e.g. Formicidae, *Canthon viridis*, etc.). Synonymous with the machine learning definition of ‘class’. |
| (Un)known & (Un)seen | Known labels are those that were input into the model. Unknown labels are those that were not input into the model. For example, a model trained using the labels “Cat”, “Dog”, and “Bird” knows the labels “Cat”, “Dog”, and “Bird” but does not know the label “Pigeon”.<br>Seen labels are those that are represented in the model via data. Seen labels may be known by the model, but it is not required. Using the same example as above, if images of pigeons were used to train the model to recognize birds, pigeons would be seen but not known by the model. If images of pigeons were not in the training dataset, pigeons would be unseen and unknown. |
| Lowest Identified Taxonomic Level (LITL) | A specimen’s LITL is the level at which the specimen’s most specific label is assigned. E.g. the LITL of a specimen labelled *Canthon* sp. is genus and it’s LITL label would be *Canthon*. |
| LITL Model | A LITL model is a machine learning model that is trained using LITL labels, usually consisting of multiple LITLs. |
| Contextual Metadata | Non-morphometric data that provides context about the time, location, and environment a specimen was collected in. |
| Zero Shot Classification | The classification of unknown and unseen specimens by making classifications at a level where the specimen’s label is known. E.g. if the beetle family Elateridae is unknown and unseen by the model, but the order Coleoptera (beetles) is known and seen by the model, Elateridae specimens can be zero shot classified at the order level. |
| Hierarchical Classification | Hierarchical classification can infer labels at higher levels from a single, base classification. These classifications can be used to calculate hierarchical accuracy at multiple levels. Zero shot classification is an example of hierarchical classification. |
| Training and Testing Datasets | The training dataset is used as the input layer of the machine learning model. It is used as the reference data to train the model how to classify each label. The testing dataset is composed of entirely separate specimens from the training dataset and is used to measure a trained model’s performance metrics (Figure S3). |
| Comprehensive Zero Shot Accuracy | Zero shot accuracy using all zero shot specimens at every taxonomic level, regardless of if they had a known taxonomic label for any given level. |
| Limited Zero Shot Accuracy | Zero shot accuracy measured only using specimens that had a known label at any given taxonomic level. |
| Top 1 Accuracy | Accuracy measured using only the label with the highest probability as measured by the model. |
| Top 3 Accuracy | Accuracy measured using the labels with the three highest probabilities as |
|  | measured by the model. If any of the three labels match the actual label, the classification is deemed correct. |
| F1 score | When referring to a model's F1 score (i.e. macro F1 score):<br>$F1 = \frac{1}{n} \sum_{i=1}^n \frac{True\ Positives_n}{True\ Positives_n + 0.5(False\ Positives_n + False\ Negatives_n)}$<br>When referring to a label's F1 score (i.e. micro F1 score):<br>$F1 = \frac{True\ Positives}{True\ Positives + 0.5(False\ Positives + False\ Negatives)}$ |
| Precision & Recall | $Precision = \frac{True\ Positives}{True\ Postivies + False\ Positives}$ $Recall = \frac{True\ Positives}{True\ Positives + False\ Negatives}$ |
| No Split | A model configuration in which no binary classifiers are used prior to the multiclass classifier. |
| 1 Split | A model configuration in which one binary classifier is used to separate intact adult specimens from everything else (e.g. body fragments, juveniles, abiotic material). The adult specimens are then passed on to a multiclass model. |
| 2 Split | A model configuration that uses the same binary classifier as the '1 Split' method, but passes the adult specimens to a second binary classifier. This classifier separates ant specimens (i.e. labels belonging to Formicidae) from non-ant specimens before passing the observations on to one of two multiclass models (an ant model or a non-ant model). |

When we trained and tested the models at the order level, XGBoost still performed the best with an average top-1 accuracy of 84.4%, top-3 accuracy of 96.1%, and F1 score of 0.683 (Figure 2). Zero-shot accuracy was 33.9% when classified to the lowest possible level. When we combined regular classifications and zero-shot classifications, the average overall accuracy of the XGBoost models across the entire NEON dataset was 73.9% when classifying at the order level or higher. The average specificity for the order level model was 4.3 (i.e. between order and class).

## Discussion

Here, we explore practical approaches for using machine learning specimen classification on a challenging terrestrial invertebrate dataset. Our dataset is continental in scope, has varying levels of taxonomic specificity, spans three phyla of terrestrial invertebrates, contains non-invertebrate and fragmented specimens, and had a very long tail distribution rank abundance curve—in short, it looks much like many real-world ecological datasets. Despite this, our best performing model (XGBoost) reached an overall LITL accuracy of 71.5% and order-level accuracy of 73.9%. These accuracies include zero-shot classifications in which the models identified taxa belonging to labels not included in the training dataset. Here we have addressed a series of common challenges in ecological datasets—taxonomic breadth, varying levels of taxonomic specificity, and uneven numbers of individuals per label—with a dataset that also has the rare feature of continental geographic range. When used in a practical setting, machine learning models will have to frequently overcome common challenges of ecological datasets while maintaining high performance standards. We propose our methods and results presented here be used as a foundation to be built and improved upon as we strive to increase the robustness and practicality of ecological machine learning models.

### The challenges and opportunities of ecological datasets

We have shown here that contextual metadata is one example of how common features of ecological datasets can be advantageous to model performance. Location data was especially important, with latitude and longitude ranking sixth and ninth out of 46 variables respectively (Figure S5). Other contextual metadata such as date, temperature, and precipitation were also more important than most morphometric variables (Figure S5). This is despite the possibility that the effect of these variables may have been dampened by the fact that no site had more than 4 sampling events and all sampling events took place in the same year. Such a small sample size means that variables like temperature and precipitation are unlikely to provide anymore information than location and time. For example, if a remote location only has one sampling event, any contextual metadata other than location would be redundant, as all specimens taken from that site would have the same contextual metadata. The fact that contextual metadata variables like temperature and precipitation still show some importance in our models testifies to their potential value in datasets spanning several years with more sampling events. In such datasets, patterns between taxa occurrence and contextual metadata could be much stronger, leading to greater variable importance and possibly higher overall accuracy.

In datasets with inconsistent taxonomic resolution, we find that LITL labelling two main advantages over single-level labelling. First, LITL labelling allows for more data to be seen by the model. In the LITL models, 96.5% of all specimens were included in either the training or testing dataset, compared to only 79.1% of specimens being included in the order-level models. This resulted in only a 1.2% difference the LITL model’s accuracy with and without zero-shot specimens (72.7% vs 71.5%), while the order-level classifier had a 10.5% gap (84.4% vs 73.9%). LITL labels also increase the model’s specificity, with our LITL models having a specificity score of 3.7 (between family and order) compared to the order-level’s specificity score of 4.3 (between order and class).

We determined that, at least in our models, taxonomic overrepresentation did not negatively impact model performance. Taxa with low prevalence in our training dataset did not have significantly lower F1 scores than taxa with high prevalence (linear regression, F = 1.568, df = 1, 42, p > 0.05, R^2^ = 0.013; Figure S6). As such, our techniques for overcoming label and taxa overrepresentation did not lead to improved performance, measured either by accuracy or F1 score (Figure 2e,f). Furthermore, in some cases, unmodified taxonomic prevalence may be beneficial, as introducing some bias to more prevalent taxa can increase accuracy. Issues surrounding bias towards high prevalent taxa arise when detecting low prevalent taxa is a higher priority than relative abundance accuracy. Models whose purpose is to measure taxonomic richness may be impacted more by prevalence bias, as low abundance taxa might be completely missed by the model. Ultimately, the importance of prevalence bias is dependant on the model’s purpose and the impact of prevalence bias on model performance.

### Challenges of measuring accuracy

Throughout the course of this study, we discovered that defining a model’s accuracy is more subjective and contextual than intuition may suggest. For our XGBoost models, accuracy can be measured as top-x (e.g. top-1, top-3, etc.), at the order-level or LITL, with zero-shot, without zero-shot, or only zero-shot. As shown in our zero-shot accuracy as well as in other studies (Blair et al., 2020), the hierarchical structure of taxonomies can also be used to measure accuracy at multiple taxonomic levels, even if the model was trained to be used at one taxonomic level. Model performance can also be measured on a per-label basis using precision, recall, and confusion matrices (Figure S7). This all leads to an overwhelming number of options to measure and interpret model performance.

To filter through this data deluge, we recommend choosing accuracy metrics that are reflective of the research question your model is addressing. For example, to estimate a model’s accuracy when it sees and knows all input taxa, an accuracy measurement that excludes zero-shot accuracy would be the most informative. Conversely, a practical measurement of accuracy that uses all possible input data, regardless of if it is seen or known by the model, including zero-shot accuracy would be best.

We also recommend researchers consider the fit of their chosen model to their research question as well. In our study, we compared our model’s order-level accuracy to the order-level accuracy of iNaturalist when predicting our specimens. While iNaturalist performed comparably to our model (84.6% vs 84.4%, respectively), there are several other factors to consider. For example, iNaturalist’s options for bulk image uploads are limited, and there is currently no way to automatically generate labels without the user manually selecting from iNaturalist’s suggestions. iNaturalist asks that no more than 10,000 lines of metadata be submitted at a time, and when we tried submitting ∼5,000 bulk images, the website became unresponsive. At the very least, a dataset like NEON’s would have to be fragmented and submitted in pieces. This might be fixable as iNaturalist’s web application is open source, so there is the possibility that enhanced bulk identification features could be added. Additionally, iNaturalist currently has no way of dealing with non-biological or fragmented specimens (i.e. specimens we listed as “Ignore”). iNaturalist assumes every submission contains at least one organism, and its model is forced to make a taxonomic classification. This can result in inaccurate classification data if images are not manually vetted for “Ignore” specimens. Finally, as a community science focused application, iNaturalist tends to be biased towards charismatic taxa. Due to this, iNaturalist performs poorly when classifying uncharismatic taxa at more specific taxonomic levels (Blair et al., 2020). Unlike the bulk classification issue, this problem is not easily solvable, as iNaturalist’s machine learning models are not open source. While iNaturalist can be an excellent tool for macroecological research, especially when used in a community science setting, other flexible options such as our models presented here are useful particularly for non-charismatic species. There is currently no one-size-fits-all solution for machine learning in an ecological context, and tools that are custom built for a particular use case are likely to be the most effective.

### Next steps

We show that zero-shot classification is highly practical for ecological purposes, as it allows otherwise unknown taxa, such as those with too few observations or with broadly specific labels to still be classified by the model. However, zero-shot classifications performed this way are still imperfect. They are far less accurate than normal classifications (33.3% less accurate in our LITL XGBoost model: 39.4% vs 72.7%), must be at a lower taxonomic specificity than their original label, and are restricted to the seen and known taxonomic groups of the model (Figure S8). This means that while the long tail of uncommon species (Figure 1) can be classified to some degree, there currently is unavoidable information loss when making these classifications. Solving this problem of unavoidable information loss and restricted labelling would be a breakthrough for zero-shot classification in ecological datasets. While it is beyond the scope of our current study, we speculate zero-shot classifications could overcome these challenges if machine learning models are paired with other classification techniques such as DNA metabarcoding. In such a tandem, the machine learning models could be informed of species presence on a per-sampling-event basis, thus increasing accuracy (by removing absent taxa from the classification pool) and specificity (by allowing species-level classifications for taxa that are only known by the model at higher levels).

### Conclusion

Automated classifications via machine learning has the potential to transform the way ecologists conduct large scale monitoring programs. However, if machine learning classification is to be accepted as a standard data collection tool for ecologists, our assessments of model performance must be demonstrative of real-life scenarios. While there is a tendency to want to be ‘fair’ to our models, fairness means very little in practice. If our models do not meet the performance demands of ecologists under field conditions, that simply means our models must be adapted and improved. Here, we demonstrated methods in which models can be assessed practically while also describing methods in which classification performance can be improved in the face of challenges posed by ecological datasets.

## Acknowledgements

This work was supported by a NSERC Discovery grant to K.E.M. as well as NSF DEB 1702426 to M.D.W., M.K., C.D.S., and K.E.M. We thank Drs Michelle Tseng, Leonid Sigal, and Rachel Germain for their fruitful discussions, as well as the National Ecological Observation Network for allowing us to use their specimens. We also thank Tanner Ortery for help in developing the imaging pipeline as part of the NSF REU program.

