## Supplemental Materials for "Embracing imperfection: machine-assisted invertebrate classification in real-world datasets"

Methods

To measure the accuracy of iNaturalist’s automatic classification tool, we submitted 13 photographs of randomly selected specimens (1 from each adult order included in our model) to iNaturalist. Accuracy was measured using iNaturalist’s first listed classification (i.e. top-1 classification). Any classifications within the correct order were deemed correct, regardless of taxonomic specificity (e.g. A species-level classification within the correct order would be recorded as correct).

Results

iNaturalist correctly classified 11 of the 13 specimens. The two orders it did not correctly classify were Opilioacarida and Zygentoma.

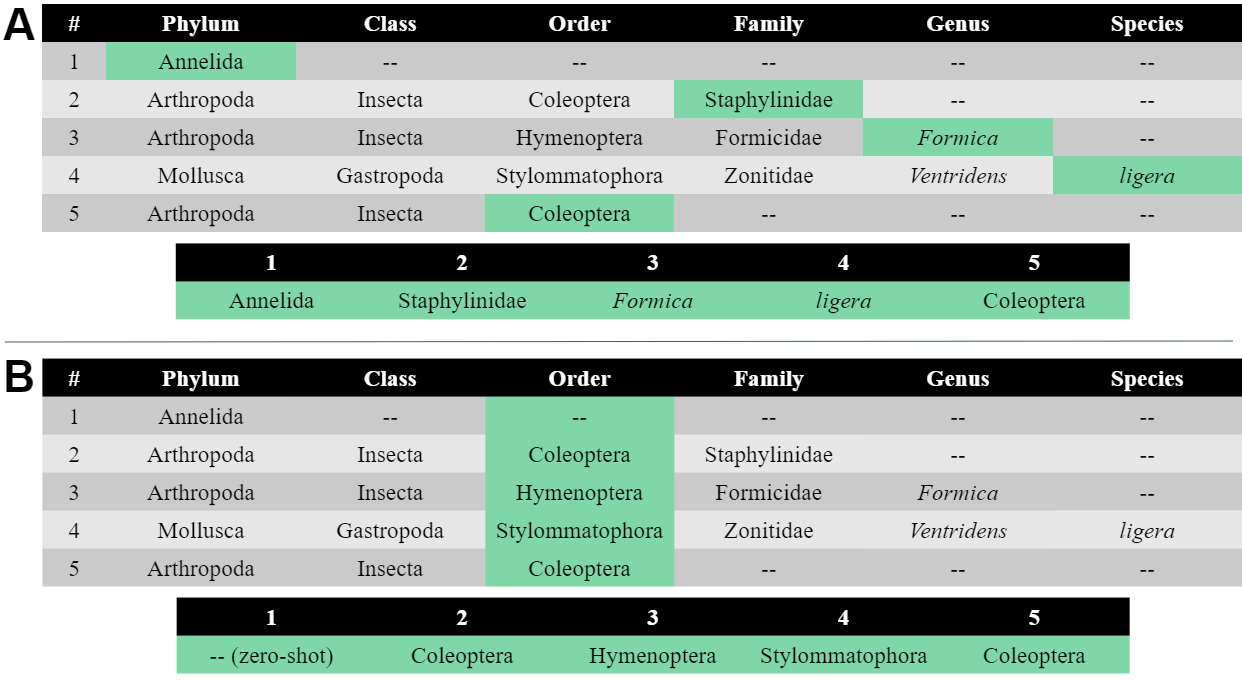

Figure S1: Lowest Identified Taxonomic Level (LITL) and order-level labelling methods. (A) The top table shows the taxonomic names (from phylum to species) for five specimens. Cells filled with “--” indicate that our labeller was unable to make a classification for that specimen at that taxonomic level. The cells highlighted in green show which label was chosen for each specimen, which are summarized in the bottom table. Labels were chosen based on the LITL for each specimen. (B) The methodology is the same as (A), except that all labels were recorded at the order-level, regardless of the LITL identified by our labeller. No order-level label was given to specimen #1, so it can only be identified using zero-shot classification.

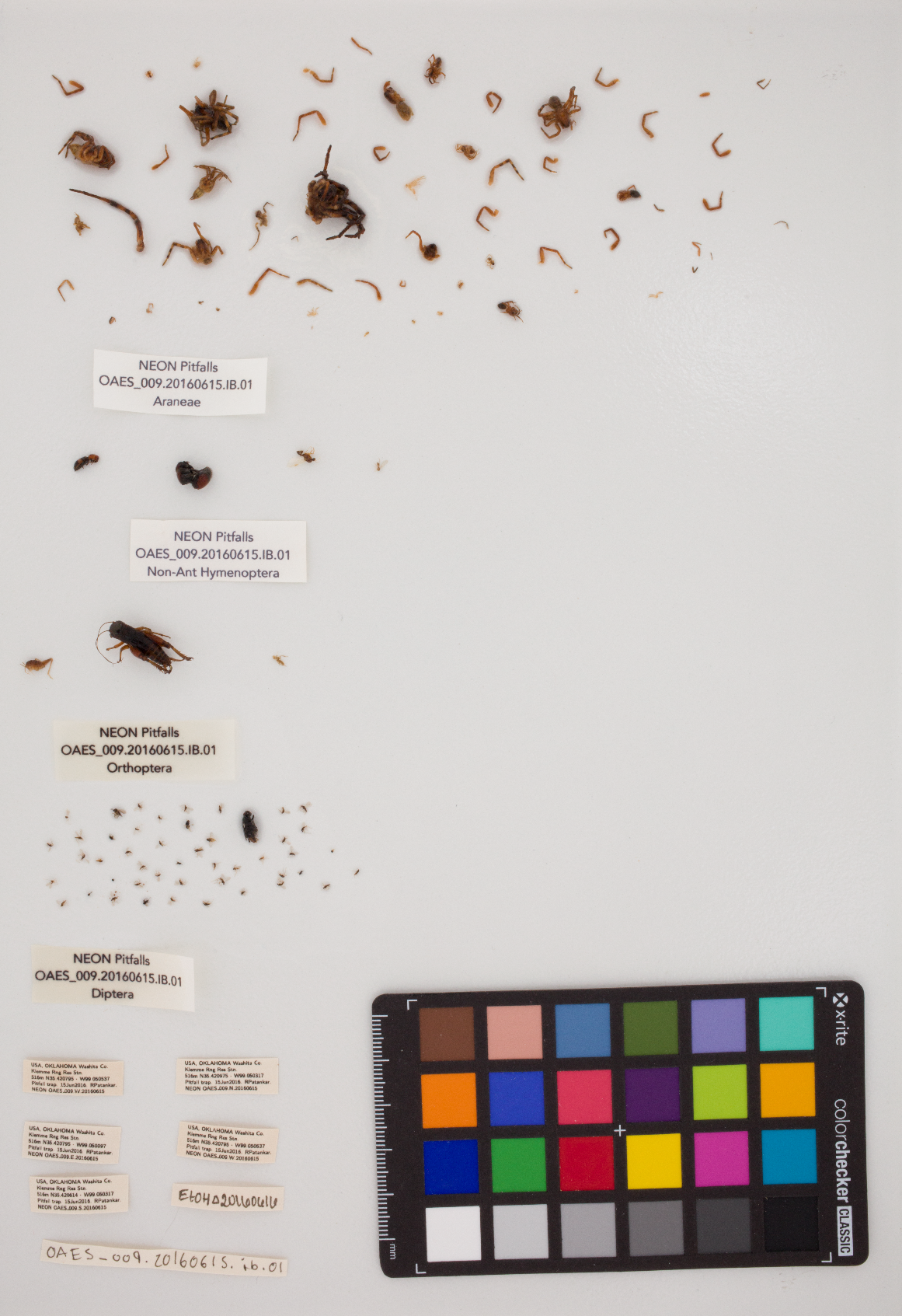

Figure S2: An example raw image of National Ecological Observatory Network (NEON) invertebrate specimens used for morphological data extraction. The specimens are clustered into groups from top to bottom: Araneae, Hymenoptera (non-ant), Orthoptera, and Diptera. It should be noted that many of the specimens in the Araneae cluster are body parts (e.g. legs) and would be labeled as ‘Ignore’ in our dataset.

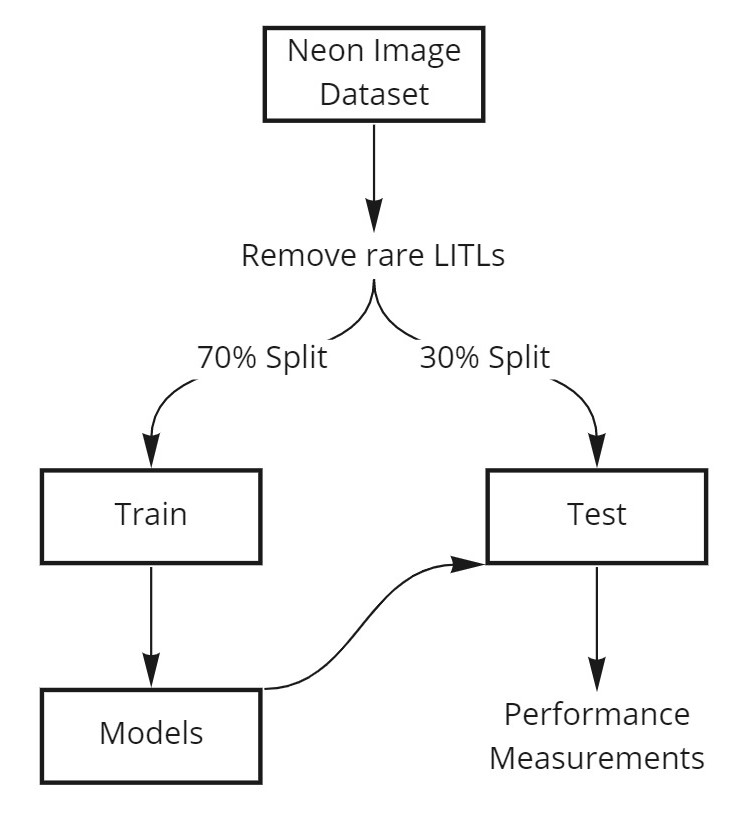

Figure S3: Flow diagram illustrating how our machine learning pipeline turns image data inputs into model performance outputs. First, LITL labels (<100 observations) are removed before the dataset is split into training and testing sets 10 times at a ratio of 70:30. Our models are trained using the training dataset, which in turn make predictions on the test data. Performance measurements such as accuracy and F1 score are measured based on the predictions made on the test datasets.

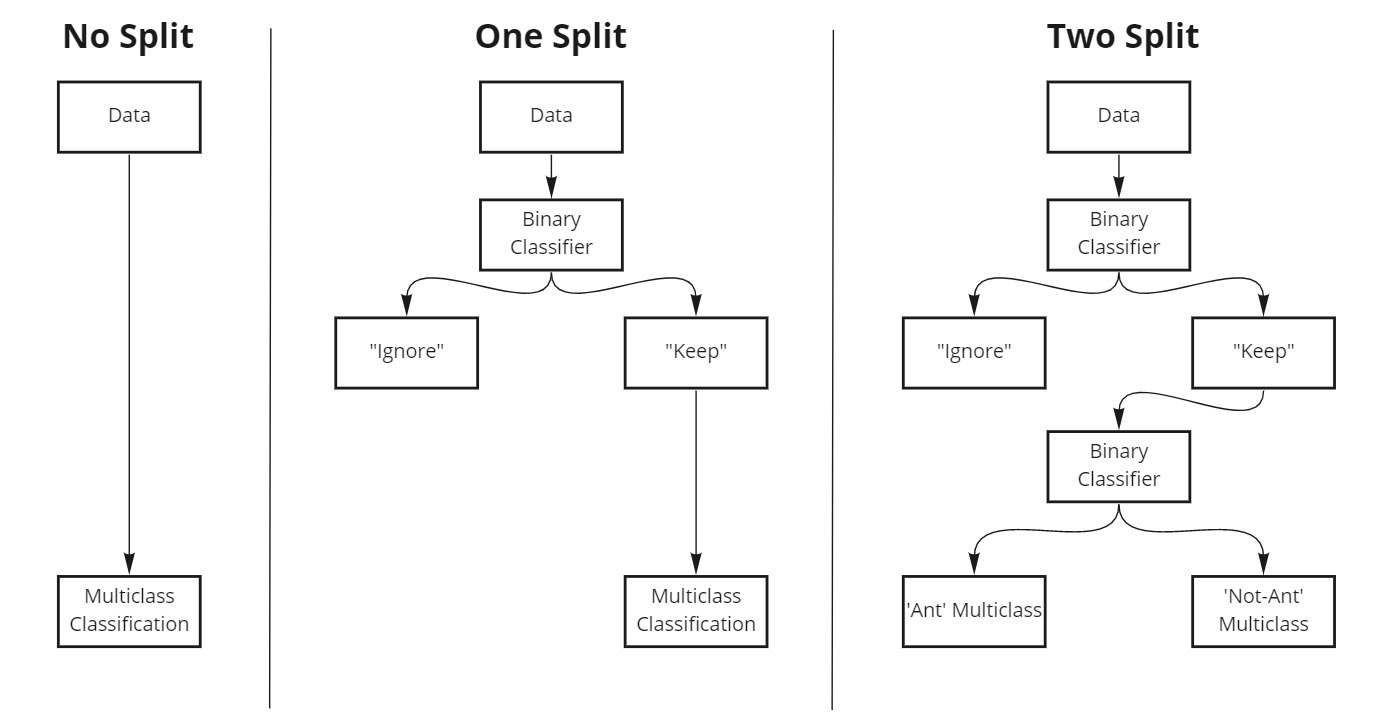

Figure S4: Binary and multiclass classifier configurations. In the “no split” configuration, the input data is put directly into a multiclass classifier with no additional modification. In the “one split” configuration, the data is first put through a binary classifier which was trained to identify “Ignore” specimens (e.g. dirt, body parts, juvenile forms, etc.). Specimens that are classified as “Keep” are passed on to a multiclass classifier for final classification, while the “Ignore” specimens are removed and labelled as “Ignore”. The “two split” configuration follows the same flow as the “one split”, but rather than passing the “Keep” specimens on to a multiclass classifier, they are passed on to a second binary classifier. This classifier labelled the specimens as either “Ants” or “Not-Ants”. Specimens labelled as “Ants” were passed on to the “Ant” multiclass classifier, which was trained only using specimens labelled as Formicidae or lower. “Not-Ant” specimens were passed on to the “Not-Ant” classifier, which was trained using all other specimens outside of Formicidae and “Ignore”.

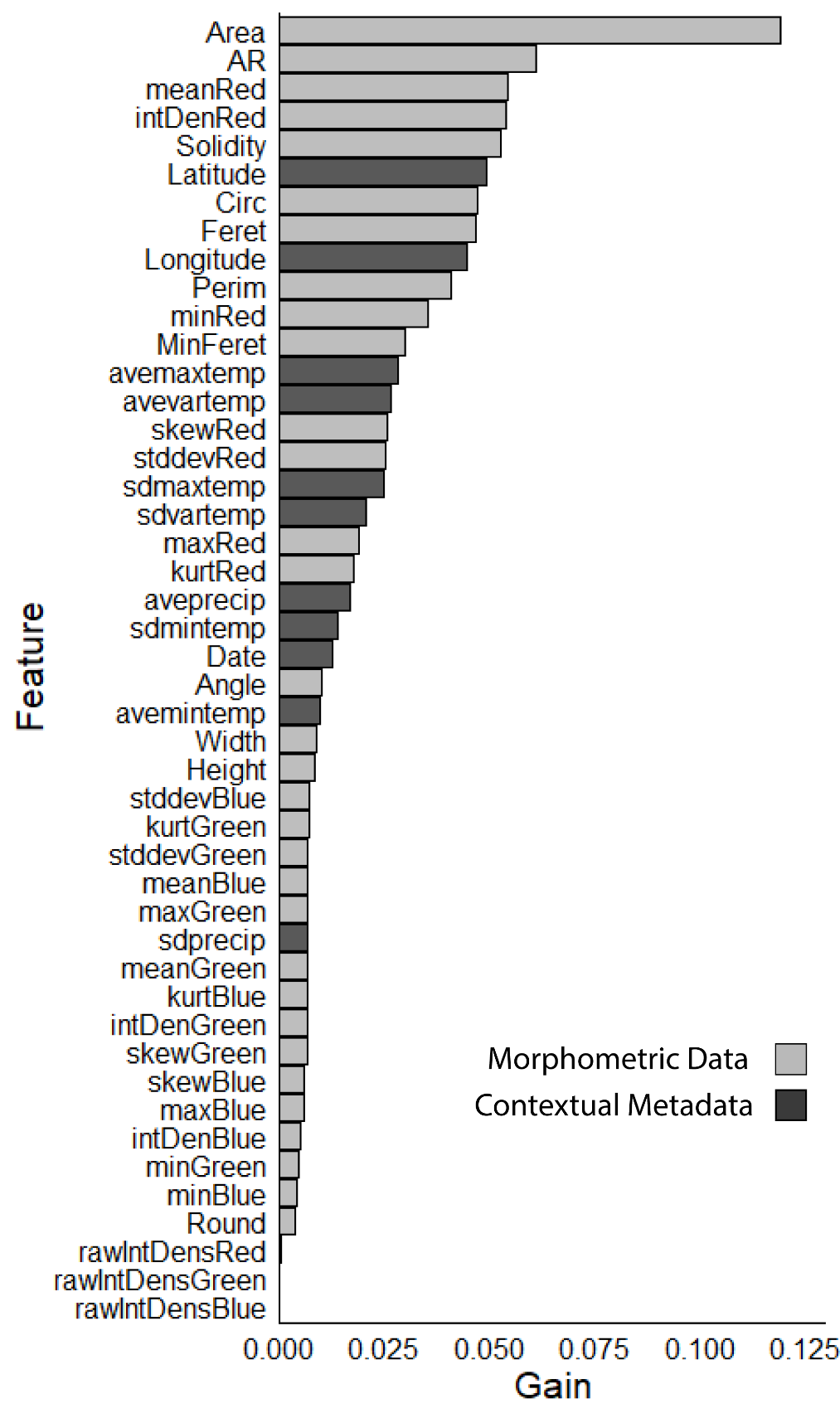

Figure S5: Predictor variable importance for the Lowest Identified Taxonomic Level (LITL) XGBoost multiclass classifier. “Gain” is defined in the xgboost R package as the contribution of each feature (predictor variable) to the model (Chen & He, 2014).

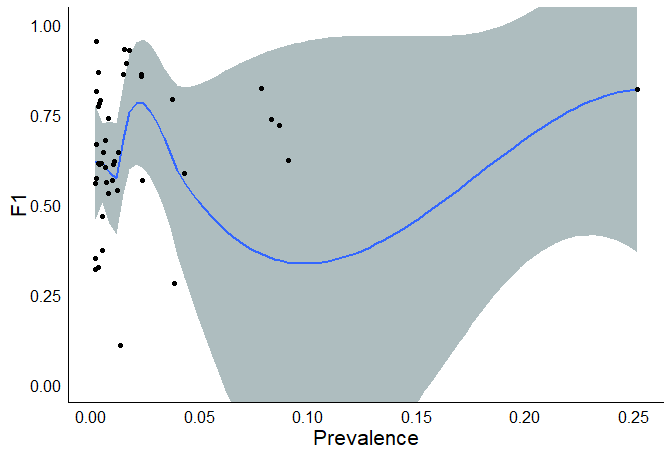

Figure S6: F1 score by prevalence for XGBoost models trained to classify invertebrate Lowest Identified Taxonomic Level (LITL) labels. Each point represents one LITL label (n = 44). A local regression curve was fit to the plot. There was no significant difference in F1 score across prevalence (F = 1.568, df = 1, 42, p > 0.05, R2 = 0.013).

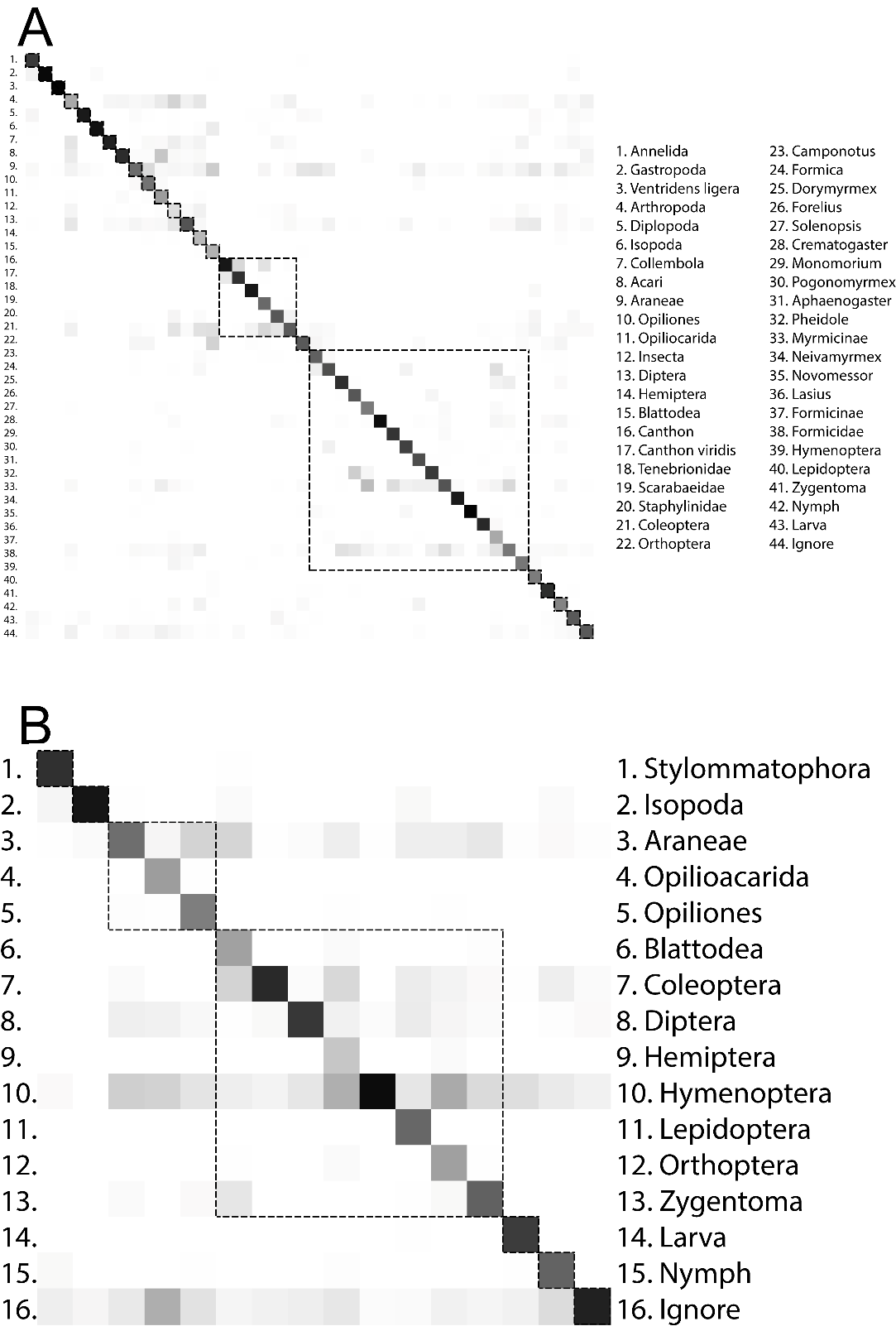

Figure S7: Confusion matrices for eXtreme Gradient Boosting (XGBoost) models trained to identify invertebrates collected by the National Ecological Observatory Network (NEON). Columns represent the actual labels while the rows represent the predicted labels. Darker boxes indicate a higher proportion of classifications within each column. The order of labels is the same between the columns and rows. (A) Lowest Identified Taxonomic Level (LITL) model confusion matrix. Dotted line boxes indicate labels belonging to the same order. (B) Order level confusion matrix. Dotted line boxes indicate labels belonging to the same class.

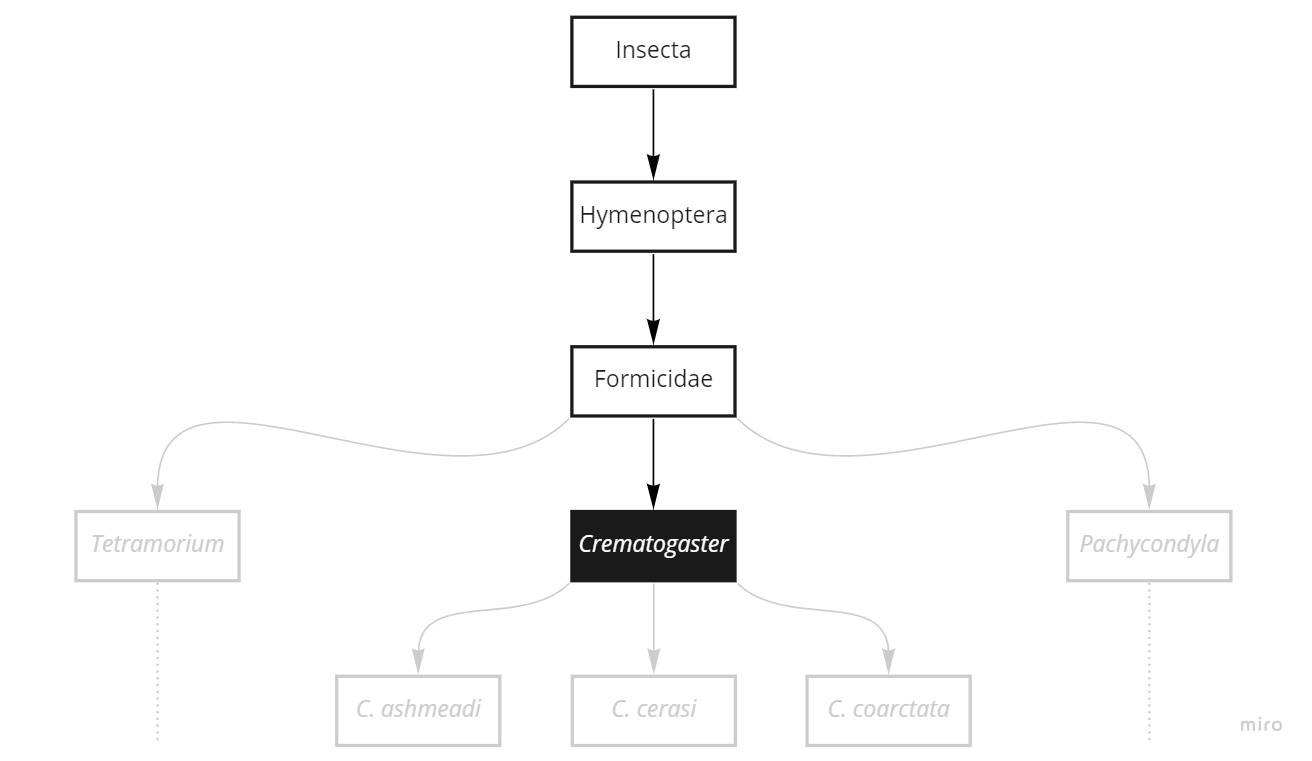

Figure S8: The possibilities and limitations of zero-shot classification. In this example, Crematogaster is known and seen by the model (shown as black box, white text), and is the label that will be output when making a classification. Using this, we can make zero-shot classifications of the parent taxa of Crematogaster (shown as black boxes with black text), as these taxa can be inferred from Crematogaster classifications. However, we cannot make zero-shot classifications for taxa that are below Crematogaster or are branches of the parental lineage as these taxa cannot be inferred from Crematogaster classifications.

Table S1: Zero-shot accuracy methods. The labels under columns Phylum to Species are the correct labels. The prediction column are the labels predicted by the model. The Known column is the label that is known by the model, and thus the first label it can possibly make a correct prediction for. Green cells indicate correct predictions, red cells are incorrect. Grey cells indicate labels the model could not possibly predict. The first results row is the ‘fair’ or ‘known’ accuracy, while the second row is the ‘unfair’ accuracy.

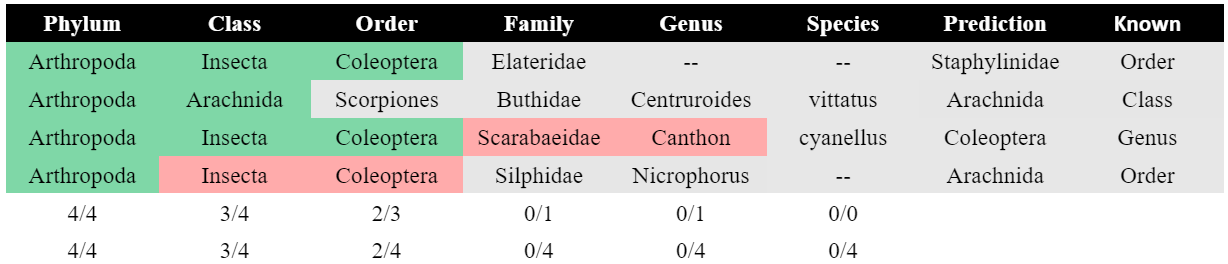
